# An *in vitro* investigation into the survival of *Tropilaelaps mercedesae* on a range of matrices

**DOI:** 10.1101/2024.12.10.627759

**Authors:** Maggie C Gill, Bajaree Chuttong, Paul Davies, Avril Earl, George Tonge, Dan Etheridge

## Abstract

*Tropilaelaps* spp. are a parasitic mite that feed and reproduce within honey bee brood (*Apis* spp.) and cause significant damage and mortality to *Apis mellifera* colonies. *T. mercedesae* is found outside the range of its natural host *A. dorsata* and was recently detected in Europe for the first time in 2024. It is widely believed that *Tropilaelaps* spp. are unable to survive without brood. However, studies have demonstrated that *T. mercedesae* can survive during broodless periods while parasitising *A. dorsata* and without brood in overwintering colonies of *A. mellifera* in temperate climates. This study sought to examine the survival of *T. mercedesae* on a range of matrices and found that mites could survive for more than 96 hours on live adult *A. mellifera* and more than 144 hours and 168 hours on decomposing *A. mellifera* pupae and adults respectively. These findings could indicate one possible mechanism which allows *T. mercedesae* to survive without brood. These findings also have implications towards better understanding possible transmission routes for *Tropilaelaps*. Previously bee sales in the form of queens and packages and used beekeeping equipment were considered a relatively low risk for transmitting *Tropilaelaps* spp. due to the absence of live brood. This study demonstrates *Tropilaelaps’* ability to survive in these scenarios and the increased potential for it to spread globally.

## 1. INTRODUCTION

Western honey bee (*Apis mellifera*) queens, packages and colonies are frequently transported on a local, national, and international level as an agricultural pollinator and for the production of bee hive products such as honey, wax and propolis. This movement of honey bees has allowed for the global transmission and spread of multiple honey bee pests and pathogens such as *Varroa destructor*, small hive beetle (*Aethina tumida*) and American foulbrood (*Paenibacillus larvae*) (Lindström, 2006; Neumann and Ellis, 2015; Roberts, et al. 2015). The ectoparasites *V. destructor* and *Tropilaelaps* spp. have jumped species from their natural honey bee hosts *Apis cerana* and *Apis dorsata* respectively to parasitise all *Apis* species of honey bee (Oldroyd, 1999; de Guzman, et al. 2016). Both mites have spread outside of the ranges of their natural hosts, with *V. destructor* now being found on every continent and in almost every country where *A. mellifera* is present (Traynor, et al. 2020; Owen, et al. 2021). Currently the spread of *Tropilaelaps* spp. is not fully verified and until recently it was believed to be restricted to Asia (Chantawannakul, et al. 2015). However, the presence of *T*. *mercedesae* has recently been confirmed in colonies of *A. mellifera* in the Krasnador and Rostov-on-don regions of Russia and the Samegrelo-Zemo Svaneti region of Georgia, thus confirming its presence in Europe for the first time (Brandorf, et al. 2024; Janashia, et al. 2024; Namin, et al. 2024).

*V. destructor* causes physical damage to honey bee brood and vectors viruses during feeding, causing significant colony losses and is considered a major threat to global beekeeping (Rosenkranz, et al. 2010). *Tropilaelaps* spp. are similar to *V. destructor* in that they primarily reproduce within sealed brood cells. Of the four know species of *Tropilaelaps* (T. *mercedesae*, *T. clareae*, *T. koenigerum* and *T. thaii*) *T*. *mercedesae* is the most widespread and is regarded as a more damaging parasite of *A. mellifera* than *V. destructor* (Anderson and Morgan, 2007). During feeding *Varroa* mites will open one or two large wounds on the brood to facilitate communal feeding of mites present within the sealed cell (Kanbar and Engles, 2005). In contrast *T. mercedesae* feed on both pre and post capped stages of brood, utilising multiple feeding sites, which can then go on to form wounds and deformities in the adult bee causing higher brood mortality than *V. destructor* infestations (Phokasem, et al. 2019). Both *V. destructor* and *T*. *mercedesae* have been shown to vector viruses when they feed (Dainat, et al. 2008; MinOo, et al. 2018). When *T*. *mercedesae* vector deformed wing virus (DWV) there is a significant reduction in the longevity and emergence weight of parasitised hosts, plus an increase to the level of DWV load and associated clinical symptoms (Khongphinitbunjong, et al. 2016). *T. mercedesae* are a major vector of honey bee pathogens and Turong, et al. (2021) demonstrated that 100% of *T. mercedesae* mites they examined harboured DWV compared to only 81.8% of *V. destructor* mites examined in the same study. Israeli acute paralysis virus (IAPV) (47.4%), sac brood virus (SBV) and *Ascosphaera apis* (31.6%) were also prevalent honey bee pathogens detected in *T. mercedesae* (Turong, et al. 2021). *T*. *mercedesae* are known to reproduce more quickly than *V. destructor* (Buawangpong, et al. 2015) and unmated females have the ability to reproduce via deuterotokous parthenogenesis, producing both male and female offspring (de Guzman, et al. 2018). The smaller size, increased mobility and shorter phoretic phase of *Tropilaelaps* spp., coupled with the similarities between the visual symptoms of infestation, makes their detection and management more difficult than that of *V. destructor* (Pettis, et al. 2013; Gill, et al. 2024). To combat the high level of colony mortality caused by *Tropilaelaps* spp. *A. mellifera* colonies kept in infested areas require continual prophylactic treatment with miticides (Rinderer, et al. 1994).

Understanding the mechanisms by which *Tropilaelaps* spp. may be transmitted is crucial to controlling their global spread. There is conflicting evidence as to whether *Tropilaelaps* spp. can survive and feed on adult bees. Rinderer, et al. (1994) reported that *Tropilaelaps* spp. did not feed on adult *A. mellifera* and were therefore unable to survive for longer than 3 days without the presence of brood. However, possible feeding of *Tropilaelaps* mites at the soft membranes of the wing axillaries was reported by de Guzman, et al. (2017). Equally under laboratory conditions Koeniger and Muzaffar (1988) observed *Tropilaelaps* survive for more than 5 days on *A. mellifera* pupae. Pettis and Chaimanee (2019) also observed *T*. *mercedesae* survive on *A. mellifera* and *A. cerana* larvae for 9-10 days and 5 days respectively under laboratory conditions. Less is known about the survival of *Tropilaelaps* spp. on their natural hosts *A. dorsata*. However, during migration *A. dorsata* colonies will ‘bivouac’ and not produce comb or brood for several weeks at a time (Robinson, 2012) with *Tropilaelaps* mites surviving by some unknown mechanism.

Beekeeping equipment and hive products pose an additional transmission route, and the World Organisation for Animal Health (OIE) recommends restricting the trade of honey bee products which are produced in colonies infested by *Tropilaelaps* mites (OIE, 2024). However, only limited research has been undertaken to assess the survival of *Tropilaelaps* mites on beekeeping equipment and hive products. Khongphinitbunjong, et al. (2019) found that *T. mercedesae* were able to survive for up to three days in dry pollen and up to six days in honeycomb and Pettis and Chaimanee (2019) found that *Tropilaelaps* mites did not survive for longer than 3 days on honey/pollen, sugar candy and royal jelly.

There are significant gaps in the existing knowledge of the survival of *Tropilaelaps* spp. and understanding their longevity and viability on hive products and other matrices has important implications to our understanding their transmissibility. This study set out to examine the survival of *T. mercedesae* on a range of matrices that represent transmission scenarios that might occur during the movement of honey bees (*A. mellifera* and *A. dorsata*), beekeeping equipment and hive products.

## 2. MATERIAL AND METHODS

The study was undertaken at Chaing Mai University, Thailand in July 2024.

### 2.1 Mite collection

Adult mites were collected from colonies located at Chiang Mai University. Brood frames from colonies where phoretic mites were observed were selected from 10 colonies. Sealed brood was uncapped using forceps and mites were collected either with an entomological aspirator (pooter) or a slightly moist fine tipped paintbrush. Mites could be encouraged to leave the brood cells by blowing over the uncapped cells or by tapping the side of the brood frame on a solid surface. Mites were visually examined and confirmed as *T. mercedesae* and pooled in a sealable plastic container (350 ml) with freshly obtained 5th instar *A. mellifera* larvae prior to being individually transferred to treatment containers.

### 2.2 Survival study

Sealable plastic containers (350 ml) were prepared with matrices and three replicates were used in each treatment group. The treatment groups comprised of

- an empty container (control),
- five *A. dorsata* adults collected from a wild colony at Chiang Mai University,
- five *A. mellifera* pupae at the pink eyed pupal stage,
- five newly emerged *A. mellifera* adults,
- five newly emerged freshly euthanised *A. mellifera* adults.

A sugar cube was glued to the side of containers containing live bees to act as a feeder and a ventilation hole covered with 0.025 mm nylon mesh was also added. *A. mellifera* pupae at the pink eyed development stage and adults were collected from frames and colonies that were randomly selected from ten colonies at Chiang Mai University. Pupae and adult bees were pooled in sealed containers before being assigned to replicate containers. Thirty *T. mercedesae* mites (n = 90 mites per treatment) were carefully individually introduced to treatment containers with a fine tipped paintbrush. Containers were then sealed with parafilm and randomly placed in an incubator at 34°C and 60% R.H. to limit possible spatial effects. *T. mercedesae* mortality was assessed every 24 hours until all the mites were dead. Mites were deemed dead if they were immobile, could not be encouraged to move when gently stimulated with a fine tipped paintbrush and were observed to have their legs curled up beneath them. It should be noted that the pink eyed pupae and dead *A. mellifera* adults were not replaced and decomposed throughout the duration of the trials.

### 2.3 Statistical analysis

All statistical analysis was performed using R Studio (R Core Team, 2018). Since the data were not normally distributed non-parametric tests were used to assess *T. mercedesae* survival between different treatments. Treatments were compared using Log Rank tests and visualized with Kaplan Meier survival curves (Fig 1.).

**Figure 1.**
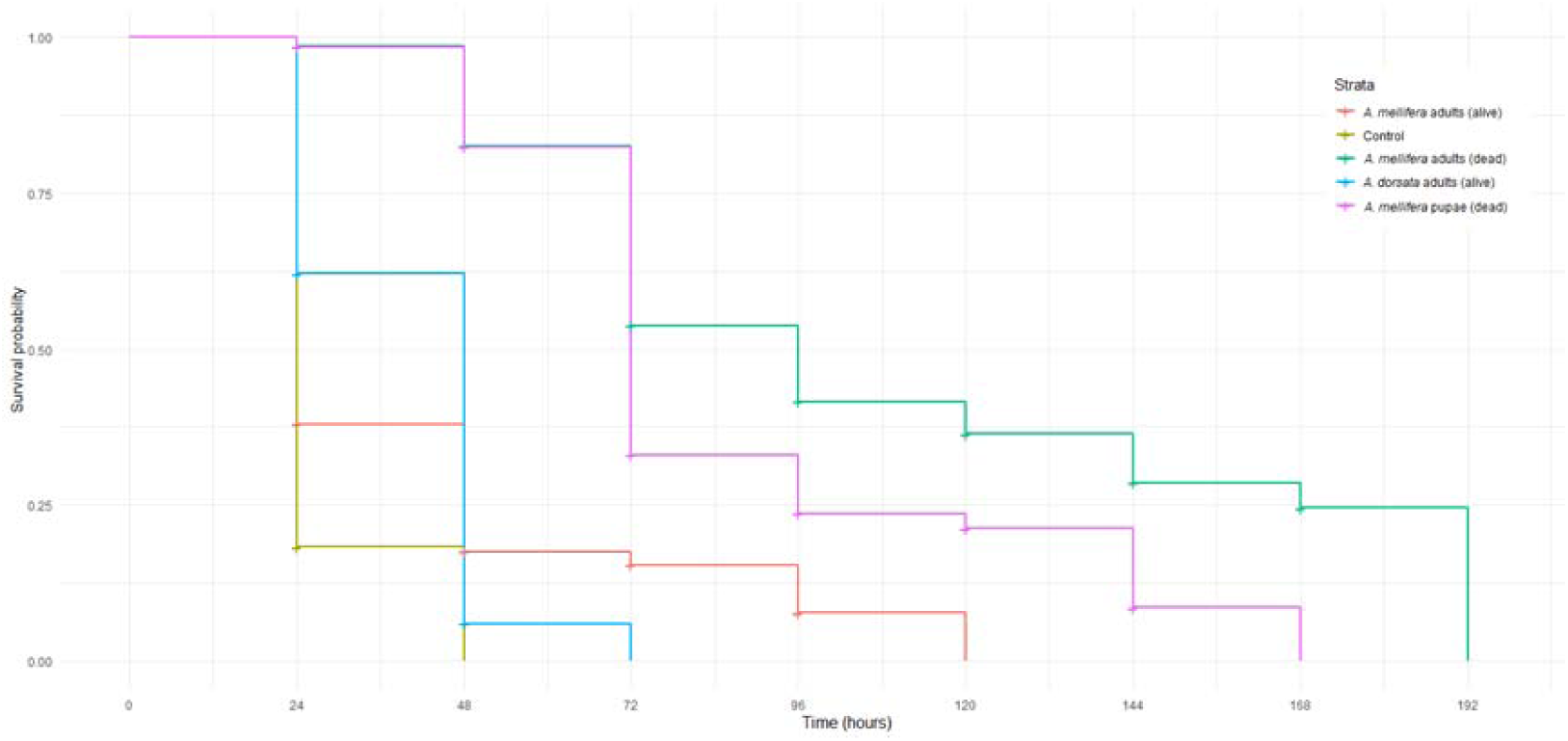
Kaplan-Meier curve displaying the survival of *T. mercedesae* mites associated with live *A. mellifera* adults, dead *A. mellifera* adults, dead *A. mellifera* pupae, live *A. dorsata* adults and the empty control treatment (*p* < 0.0001) (n = 30 mites / treatment).

## 3. RESULTS

The mean survival time of *T. mercedesae* mites was 13.3 hours in the control treatment, 15 hours in the treatment group containing live *A. mellifera* adults, 21.8 hours in the treatment group containing live *A. dorsata* adults, 54.6 hours in the treatment group containing dead *A. mellifera* pupae and 55.6 hours in the treatment group containing dead *A. mellifera* adults. The final *T. mercedesae* died after 168 hours in the treatment group containing dead *A. mellifera* adults and survived for significantly longer than mites in the other treatment groups (Fig 2.). A log rank test (Wilcoxon - Breslow) determined that the survival distributions between the differing treatments were statistically significantly (χ^2^(4) = 478.71, *p* <0.0001). Pairwise comparison using the Log Rank test showed that there was a statistically significant difference in survival distributions between all treatment groups apart from live *A. dorsata* adults vs live *A. mellifera* adults and *p* values are shown in Table 1.

**Figure 2.**
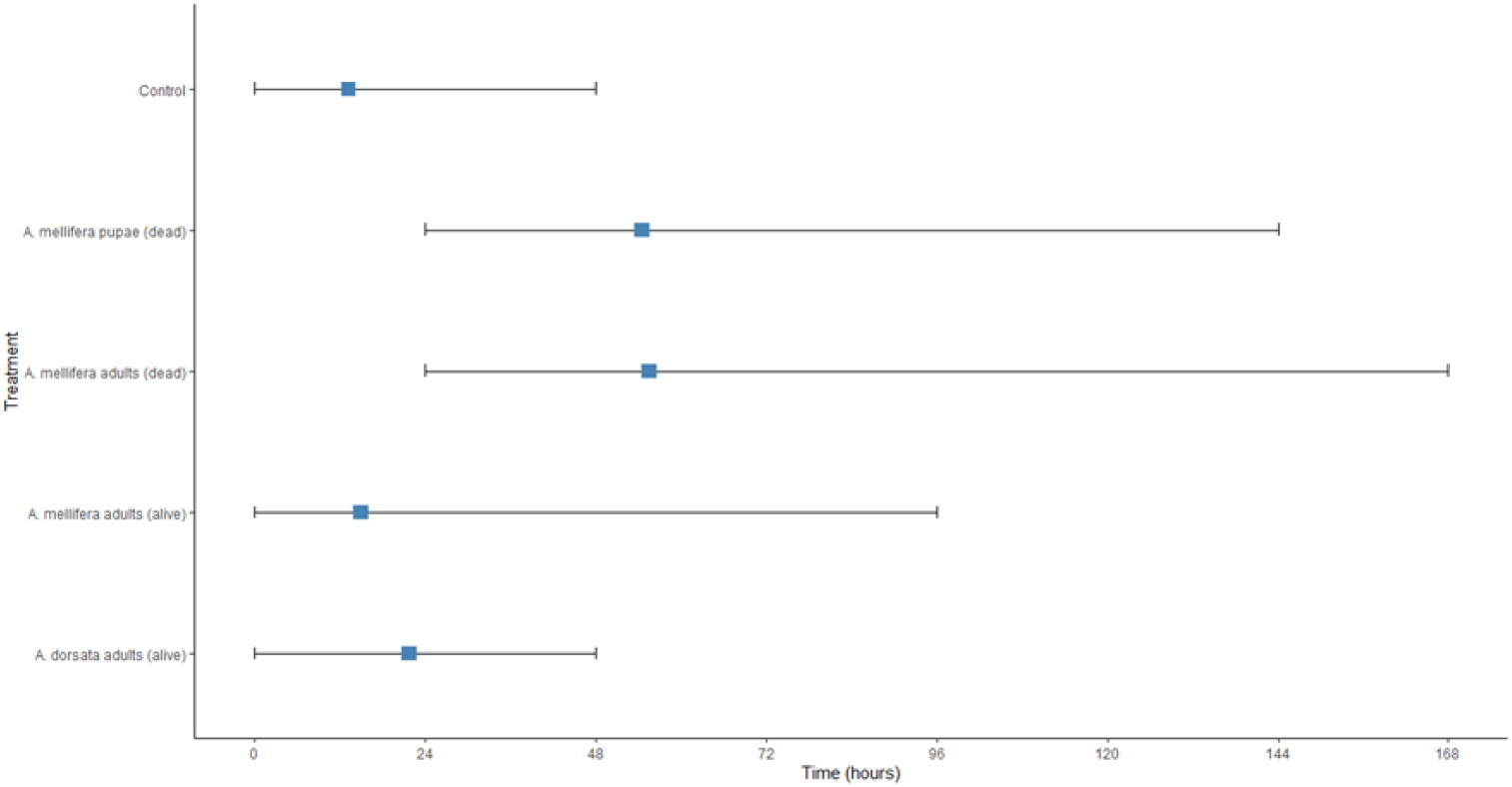
Diagram showing the survival times of *T. mercedesae* with blue squares showing mean survival time. All data that support the findings of this study are included within this paper and its Supplementary Information files.

**Table 1.** *p* values from pairwise comparison using the Log Rank test.

|  | Live *A. mellifera* adults | Control | Dead *A. mellifera* adults | Live *A. dorsata* adults |
| --- | --- | --- | --- | --- |
| Control | $p < 0.001$ | - | - | - |
| Dead *A. mellifera* adults | $p < 0.0001$ | $p < 0.0001$ | - | - |
| Live *A. dorsata* adults | $p = 0.0776$ | $p < 0.0001$ | $p < 0.0001$ | - |
| Dead *A. mellifera* pupae | $p < 0.0001$ | $p < 0.0001$ | $p < 0.01$ | $p < 0.0001$ |

## 4. DISCUSSION

*T. mercedesae* mites experienced high mortality in the control group, with 90% of mites not surviving for more than 24 hours in the treatment which consisted of an empty container with no food source, and these results corroborate those of Koeniger and Muzaffar, (1988) and Pettis and Chaimanee, (2019). However, *T. mercedesae* survived for longer than previously observed in other studies in the treatment groups containing dead *A. mellifera* adults, dead *A. mellifera* pupae and live *A. mellifera* adults (Koeniger and Muzaffar, 1988; Pettis and Chaimanee, 2019). De Guzman, et al. (2017) observed the apparent feeding of *T. mercedesae* on adult *A. mellifera* where the mites mouthparts appeared to pierce the soft membrane of the wing axillaries and this was accompanied by a pumping or pulsating motion of the opisthosoma. Feeding on live adult bees was not observed during this study, however a small proportion of mites (∼10%) were able to survive for significantly longer on live adult *A. mellifera* than mites in the control group, which could suggest that these mites were able to feed on the adult bees. Another possibility is that *Tropilaelaps,* like *Braula coeca,* may receive food during trophallaxis between bees. Koeniger and Muzaffar (1988) observed the “conspicuous reactions of mites” during trophallaxis between *A. dorsata* and suggested that mite feeding might be taking place. This theory might also explain why the survival distribution between the live *A. mellifera* and live *A. dorsata* treatments was not statistically significantly different, although both groups did survive for significantly longer than the control group. Mites did not survive for longer than 48 hours in the treatment group containing live *A. dorsata* adults, which is again in line with the findings of Koeniger and Muzaffar (1988). Koeniger and Muzaffar (1988) observed injuries and lost appendages on *T. mercedesae* in their *A. dorsata* treatment groups, whereas the mites in other treatment groups were uninjured. The dead mites in this study were not examined for injuries, however *T. mercedesae* survived for up to 48 hours longer on live *A. mellifera* when compared to live *A. dorsata* and this may be due in part to the parasite host relationship and grooming behaviours of *A. dorsata*.

*T*. *mercedesae* mites survived for up to 168 hours in the treatment group containing dead *A. mellifera* pupae and up to 192 hours in the treatment group containing dead *A. mellifera* adults and it should be noted that dead pupae and adults were not replaced throughout the trial but instead decomposed. It is unclear if Pettis and Chaimanee (2019) replenished the *A. mellifera* larvae used in their study to achieve mite survival of 9-10 days (216 – 240 hours) or allowed them to decompose, however Koeniger and Muzaffar (1988) replaced *A. mellifera* pupae every 3 days and observed mite survival of up to 120 hours. Mites were observed spending the majority of their time on the dead pupae and dead bees and appeared to feed on the exudate created during decomposition. This finding suggests that *T. mercedesae* could be transported in scenarios where live *A. mellifera* brood and adults are not present, such as in used beekeeping equipment containing decaying brood and bees or in queen shipments and packages where bees have died during the caging / packaging and transportation process. Feral colonies have been transported to countries outside of the current distribution of *T. mercedesae* on shipping containers, boats, and airplanes (Heersink, et al. 2016) and it has been assumed that broodless colonies and colonies that have died at sea do not pose a transmission risk for *Tropilaelaps* spp. (ESFA, 2013). Our findings suggest that aside from the obvious transmission root of brood within feral *Apis* spp. colonies *T. mercedesae* could survive in a broodless scenario on live adult bees or on decaying brood or adult bees that have died during transportation.

*T. mercedesae* can survive during broodless periods in *A. mellifera* colonies (Brandorf, et al. 2024) and during broodless periods on their natural hosts *A. dorsata* during bivouacking (Robinson, 2013), but the mechanisms by which they are able to do so are little researched and not understood. This study has demonstrated *T. mercedesae* ability to survive on decaying *A. mellifera* brood and bees and live newly emerged *A. mellifera* bees, and these findings coupled with that of Robinson (2013) and Brandorf, et al. (2024) demonstrate that the transmissibility of *T. mercedesae* is higher than previously evaluated. Given the increasing global spread of *T. mercedesae* and its recent incursion into regions bordering areas which produce significant amounts of honey each year such as Ukraine and Türkiye (World Population Review, 2024) and given that political and socioeconomic factors in these regions could impact successful monitoring for *Tropilaelaps* spp. it is important that beekeepers and authorities globally are vigilant to the threat that the transmission of *Tropilaelaps* poses.

## Supporting information

https://10.6084/m9.figshare.28001570

## ACKNOWLEDGEMENT.

This work was funded with a grant from Bee Disease Insurance ltd (https://www.beediseasesinsurance.co.uk/) and a donation from The Worshipful Company of Wax Chandlers (https://www.waxchandlers.org.uk/). The authors thank Victoria Tomkies for advice on experimental design and help with the preliminary study.

## REFERENCES.

Anderson DL, Morgan MJ. (2007). Genetic and morpho-logical variation of bee parasitic *Tropilaelaps* mites (Acari: Laelapidae): New and re-defined species. Exp Appl Acarol. 43: 1–24. DOI:10.1007/s10493-007-9103-0

Brandorf A, Ivoilova MM, Yanez O, Neumann P, Soroker V. (2024) First report of established mite populations, *Tropilaelaps mercedesae*, in Europe. J Apic Res. DOI: 10.1080/00218839.2024.2343976

Buawangpong N, de Guzman LI, Khongphinitbunjong K, Frake AM, Burgett M, Chantawannakul P. (2015). Prevalence and reproduction of *Tropilaelaps mercedesae* and *Varroa destructor* in concurrently infested *Apis mellifera* colonies. Apidologie. 46:779–86. DOI: 10.1007/s13592-015-0368-8

Chantawannakul P, de Guzman LI, Li J, Williams GR. (2015). Parasites, pathogens, and pests of honeybees in Asia. Apidologie. 47:301–24. DOI: 10.1007/s13592-015-0407-5

Dainat B, Ken T, Berthoud, H. Neumann P. (2008). The ectoparasitic mite *Tropilaelaps mercedesae* (Acari, Laelapidae) as a vector of honeybee viruses. Insect. Soc. 56: 40–43. DOI; 10.1007/s00040-008-1030-5

de Guzman LI, Williams GR, Khongphinitbunjong K, Chantawannakul P. (2017) Ecology, Life History, and Management of *Tropilaelaps* Mites, J Econ Entomol. 110(2): 319–332. DOI: 10.1093/jee/tow304

de Guzman LI, Phokasem P, Khongphinitbunjong K, Frake, AM, Chantawannakul P. (2018) Successful reproduction of unmated *Tropilaelaps mercedesae* and its implication on mite population growth in *Apis mellifera* colonies. J. Invertebr. Pathol. 153: 35–3. DOI: 10.1016/j.jip.2018.02.010

EFSA Panel on Animal Health and Welfare (AHAW). (2013). Scientific Opinion on the risk of entry of *Aethina tumida* and *Tropilaelaps* spp. in the EU. EFSA Journal. 11(3):3128.

Gill MC, Chuttong B, Davies P, Etheridge D, Panyaraksa L, Tomkies V, Tonge G, Budge GE. (2024). Assessment of the efficacy of field and laboratory methods for the detection of *Tropilaelaps* spp. PLoS One. 19(9). DOI: 10.1371/journal.pone.0301880

Heersink DK, Caley P, Paini DR, Barry SC. (2016). Quantifying the establishment likelihood of invasive alien species introductions through ports with application to honeybees in Australia. Risk Analysis. 36(5):892–903. DOI: https://onlinelibrary.wiley.com/doi/epdf/10.1111/risa.12476?saml_referrer

Janashia I, Uzunov A, Chen C, Costa C, Cilia G. (2024). First report on *Tropilaelaps mercedesae* presence in Georgia: The mite is heading westward! J Api Sci. Sci. 0: 10.2478/jas-2024-0010

Kanbar G. Engels W. (2005). Communal use of integumental wounds in honey bee (*Apis mellifera*) pupae multiply infested by the ectoparasitic mite *Varroa destructor*. Genet Mol Res. 4: 465–472

Khongphinitbunjong K, Neumann P, Chantawannakul P, Williams GR. (2016) The ectoparasitic mite *Tropilaelaps mercedesae* reduces western honey bee, Apis mellifera, longevity and emergence weight, and promotes Deformed wing virus infections. J Invertebr Patholo. 1(137):38–42.

Khongphinitbunjong K, Chantawannakul P, Yañez O, Neumann P. (2019). Survival of ectoparasitic mites *Tropilaelaps mercedesae* in association with honeybee hive products. Insects. 10(2):36.

Koeniger N, Muzaffar N. (1988). Life-span of the parasitic honeybee mite, *Tropilaelaps clareae*, on *Apis cerana*, *Apis dorsata* and *Apis mellifera*. J. Apic. Res. 27: 207– 212

Lindström A. (2006) Distribution and transmission of American foulbrood in honey bees. No. 2006: 22.

MinOo H, Kanjanaprachoat P, Suppasat T, Wongsiri S. (2018) Honey bee virus detection on *Tropilaelaps* and *Varroa* mites in Chiang Mai Thailand. J Apiculture. 33(2):77–81. DOI: 10.17519/apiculture.2018.06.33.2.77

Namin SM, Joharchi O, Aryal S, Thapa R, Kwon S, Kakhramanov BA, et al. (2024) Exploring genetic variation and phylogenetic patterns of *Tropilaelaps mercedesae* (Mesostigmata: Laelapidae) populations in Asia. Front. Ecol. Evol. 12: DOI: 10.3389/fevo.2024.1275995

Neumann P, Ellis JD. (2008) The small hive beetle (*Aethina tumida* Murray, Coleoptera: Nitidulidae): distribution, biology and control of an invasive species. J Apicult Res. 47(3): 181–183. DOI: 10.1080/00218839.2008.11101453

OIE. Terrestrial Animal Health Code. (2024). Section 9. Apinae. Chapter 9.5. Infestation of Honeybees with *Tropilaelaps* spp. DOI: https://www.woah.org/fileadmin/Home/eng/Health_standards/tahc/2023/chapitre_tropilaelaps_spp.pdf

Oldroyd, BP. (1999) Coevolution while you wait: *Varroa jacobsoni*, a new parasite of western honeybees. Trends Ecol Evol. 14: 312–315

Owen R, Stevenson M, Scheerlinck JP. (2021). *Varroa destructor* detection in non-endemic areas. Apidologie. 52:900–14. DOI: 10.1007/s13592-021-00873-7

Pettis JS, Rose R, Lichtenberg EM, Chantawannakul P, et al. (2013). A rapid survey technique for *Tropilaelaps* mite (Mesostigmata: Laelapidae) detection. J Econ Entomol. 106(4): 1535–1544.

Pettis JS, Chaimanee V. (2019). The survival of *Tropilaelaps mercedesae* on beehive products. J Apicult Res. 58(3): 413–5.

Phokasem P, de Guzman LI, Khongphinitbunjong K. Frake AM. Chantawannakul P. (2019) Feeding by *Tropilaelaps mercedesae* on pre- and post-capped brood increases damage to *Apis mellifera* colonies. Sci Rep. 9. DOI: 10.1038/s41598-019-49662-4

R Core Team. (2018). R: A Language and Environment for Statistical Computing. R Foundation for Statistical Computing. R Core Team: Vienna, Austria.

Rinderer TE, Oldroyd BP, Lekprayoon C, Wongsiri S, et al. (1994) Extended survival of the parasitic honey bee mite *Tropilaelaps clareae* on adult workers of *Apis mellifera* and *Apis dorsata*. J Apicult Res. 33(3): 171–174.

Roberts JM, Anderson DL, Tay WT. (2015) Multiple host shifts by the emerging honeybee parasite, *Varroa jacobsoni*. Mol Ecol. 24(10): 2379–91. DOI: 10.1111/mec.13185.

Robinson WS. (2012). Migrating giant honey bees (*Apis dorsata*) congregate annually at stopover site in Thailand. PLoS One. DOI: 10.1371/journal.pone.0044976

Rosenkranz P, Aumeier P, Ziegelmann B. (2010). Biology and control of *Varroa destructor*. J Invertebr Patholo. 103: S96–S119. DOI: 10.1016/j.jip.2009.07.016

Truong AT, Yoo MS, Yun BR, Kang JE, Noh J, Hwang TJ, Seo SK, Yoon SS, Cho YS. (2021). Prevalence and pathogen detection of *Varroa* and *Tropilaelaps* mites in *Apis mellifera* (Hymenoptera, Apidae) apiaries in South Korea. J Apicult Res. 8;62(4):804-12.

Traynor KS, Mondet F, de Miranda JR, Techer M, Kowallik V, et al. (2020) *Varroa destructor*: A complex parasite, crippling honey bees worldwide. Trends parasitol. 36(7): 592–606.

World Population Review. (2024) Honey production by country 2024. Available from https://worldpopulationreview.com/country-rankings/honey-production-by-country

